# FlexDotPlot: a universal and modular dot plot visualization tool for complex multifaceted data

**DOI:** 10.1101/2020.04.03.023655

**Authors:** Simon Leonard, Aurélie Lardenois, Karin Tarte, Antoine Rolland, Frédéric Chalmel

## Abstract

**Motivation:** Dot plots are heatmap-like charts that provide a compact way to simultaneously display two quantitative information by means of dots of different sizes and colours. Despite the popularity of this visualization method, particularly in single-cell RNA-seq studies, existing tools used to make dot plots are limited in terms of functionality and usability.

**Results:** We developed FlexDotPlot, an R package for generating dot plots from multifaceted data, including single-cell RNA-seq data. It provides a universal and easy-to-use solution with a high versatility. An interactive R Shiny application is also available allowing non-R users to easily generate dot plots with several tunable parameters.

**Availability and implementation:** Source code, detailed manual, and code to reproduce figures are available at https://github.com/Simon-Leonard/FlexDotPlot. A Shiny app is available as a stand-alone application within the package.

## Introduction

Data visualization is essential for the biological mining of the vast amount of information generated by high-throughput technologies. One of the most popular plotting technique in genomics is the heatmap. It is used to display a single quantitative information from a two-dimensional table (Wilkinson and Friendly, 2009). For instance, in transcriptomics, rows and columns usually represent genes and samples, respectively, and boxes are colour-coded according to expression signals. Heatmaps however appear intrin-sically limited to provide comprehensive representation of the diversity of information now available from single-cell technolo-gies. Accordingly, alternative methods such as dot (or spot) plots are increasingly used.

A dot plot is a modified heatmap where each box in the grid is replaced by a dot. In addition to quantitative values that are dis-played via a colour gradient, the dot size is also used to represent another quantitative information, e.g. the fraction of cells express-ing a given gene within a single-cell RNA-sequencing (scRNA-seq) dataset (Lukassen et al., 2018; Habib et al., 2017; Ordovas-Montanes et al., 2018; Wu et al., 2018).

In recent years, multiple tools have been developed to make dot plots like the rain plot method (Henglin et al., 2019) or the corrplot function of the feature-expression heatmap method (Benno Haar-man et al., 2015; Wei and Simko, 2017). Several programs dedi-cated to scRNA-seq analysis (Seurat, scClustViz or cellphonedb) also provide a dot plot function (Innes and Bader, 2019; Stuart et al., 2019; Efremova et al., 2019). A dot plot generator is also available in ProHits-viz, a web-tool dedicated to protein-protein interaction analysis (Knight et al., 2017).

While the increasing number of dedicated tools embedding dot plot representation clearly illustrates the relevance of these visualiza-tion methods in current data analysis, dot plots remain difficult to generate partly due to the lack of universality. In particular, most of the previously mentioned packages use very different input data formats: rain plot and corrplot are restricted to correlation and association matrixes respectively; cellphonedb, Seurat and scClustViz require an output from their own pipeline with a specif-ic format; ProHits-viz requires quantitative information on bait-prey interaction. In addition, all of the existing methods can only display two quantitative information per dot (Supplementary Table 1). To address these issues, we developed a simple and universal tool for dot plot visualization.

## Implementation

FlexDotPlot is implemented in R and takes advantage of several publicly available R packages for data visualization, manipulation and analysis (Supplementary Table 2). It requires a simple data frame as input: the first two columns contain the two factors to spread along the x and y axes (e.g. genes and cell populations for scRNA-seq datasets), followed by the corresponding continuous and/or discrete data to be displayed (Supplementary Fig. S1A). From this input, dot plots can be produced with a single command line or interactively with a Shiny application.

## Features highlights

FlexDotPlot consists of a single function to generate dot plots with several easy-to-tune parameters allowing users to specify which and how information has to be represented (Supplementary Fig. S1B). In addition to the traditional size and colour features, users can display two additional information by adding some text or by using dot shapes (Fig. 1).

**Figure 1:**
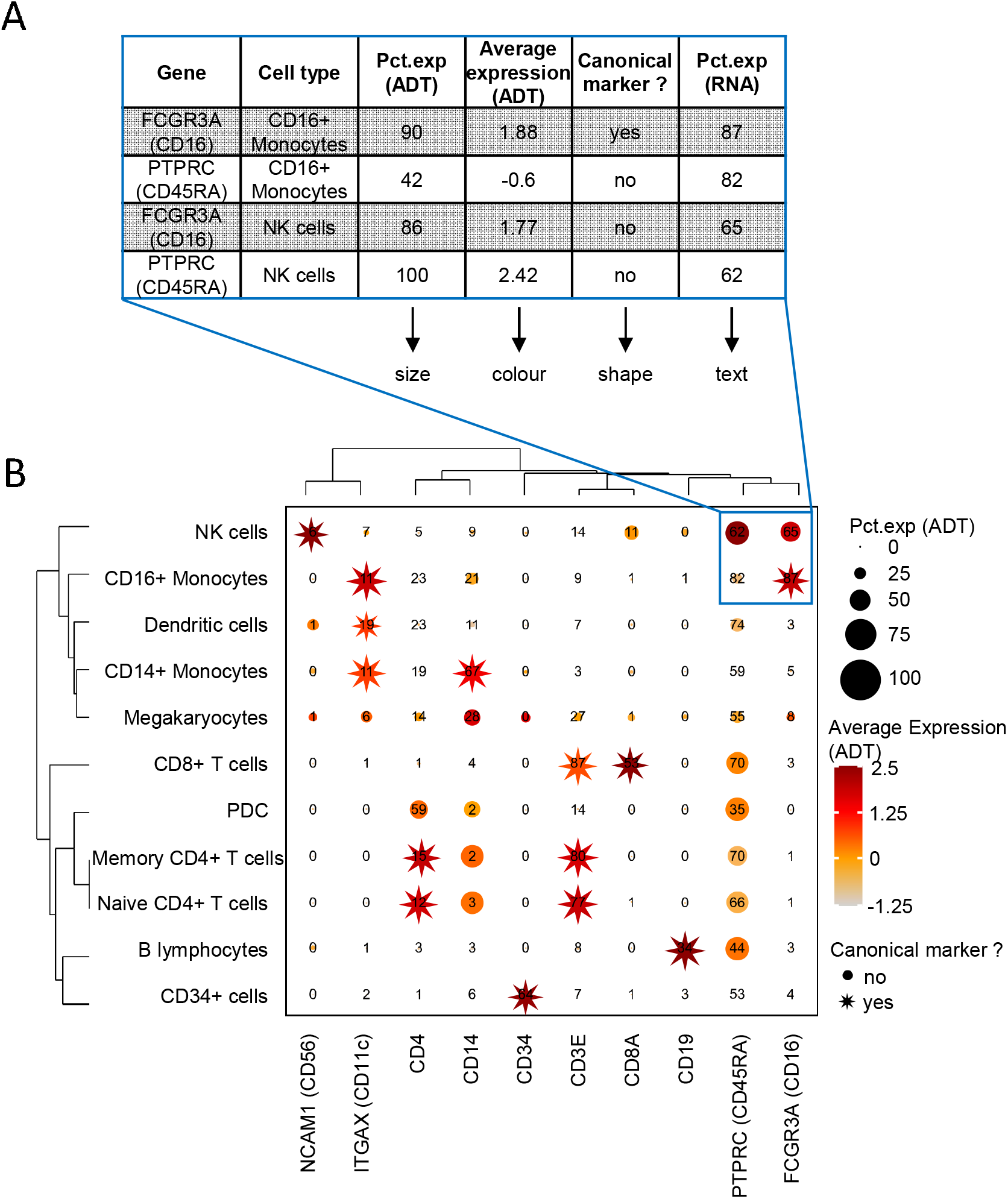
DotPlot representation applied to the 8k human CBMC scRNA-seq dataset. **A.** Part of the input table used for generating dotplot in B. **B.** Dotplot on the CBMC dataset with FlexDotPlot. Each dot represents multiple features for one gene (vertical axis) in a given cell cluster (horizontal axis). The size and text of each dot represent the percentage of cells exprsseing a given marker (Pct.exp) according to Antibody Derived Tags (ADT) and RNA level respectively. The colour represents the average scaled expression of a given marker at the ADT level. The shape of the dot is used to highlight canonical marker of specific cell types. Blue border highlights the part of the plot corresponding to the table in A. **Data source:** SeuratData package (https://github.com/satijalab/seurat-data)

One major improvement of FlexDotPlot relies on the possibility to represent a defined factor by using variable dot shapes, which is not possible with other existing dot plot representation methods (Supplementary Table 1). This characteristic is really suitable to represent percentages as it is classically done in scRNA-seq related dot plots (Supplementary Fig. S2).

Several minor parameters are implemented in the function to easily custom the plot legends, shapes, colours and/or labels. Moreover, a ggplot2 object can be returned so that users can customize even further the output by adding custom layers if needed.

FlexDotPlot also provides parameters to enhance the resulting figure with dendrograms, which is rarely available in existing methods (Supplementary Table 1) (Wilk et al., 2020). Depending on the variable types, a principal component analysis (continuous variables), a multiple correspondence analysis (discrete variables) or a factor analysis for mixed data (continuous and discrete varia-bles) is performed prior to clustering analysis.

To illustrate further the versatility and usefulness of FlexDotPlot in representing multimodal data, we used FlexDotPlot on several scRNA-Seq datasets combined with complementary results including CITE-seq, CNV or cell communication data (Stoeckius et al., 2017; Bi et al., 2021; Tirosh et al., 2016). Particularly, we repro-duced dot plot figures from published articles or tutorials and proposed enhanced versions using FlexDotPlot (Supplementary Fig. S2, S3, S4).

## Conclusion

FlexDotPlot is an easy to use tool for generating highly customizable dot plot representations. It uses standardized input and output formats (data frame and *ggplot2* object respectively), which also makes the combination of FlexDotPlot and other R pipelines possible. A set of vignettes are included in the package to illustrate the dot plot construction procedures and their associated parameters.

## Acknowledgments

We thank Paul Rivaud for his critical reading and members of the GenOuest BioInformatics core facility for their support

