## Supplementary Figures for "FlexDotPlot: a universal and modular dot plot visualization tool for complex multifaceted data"

### Slide 1
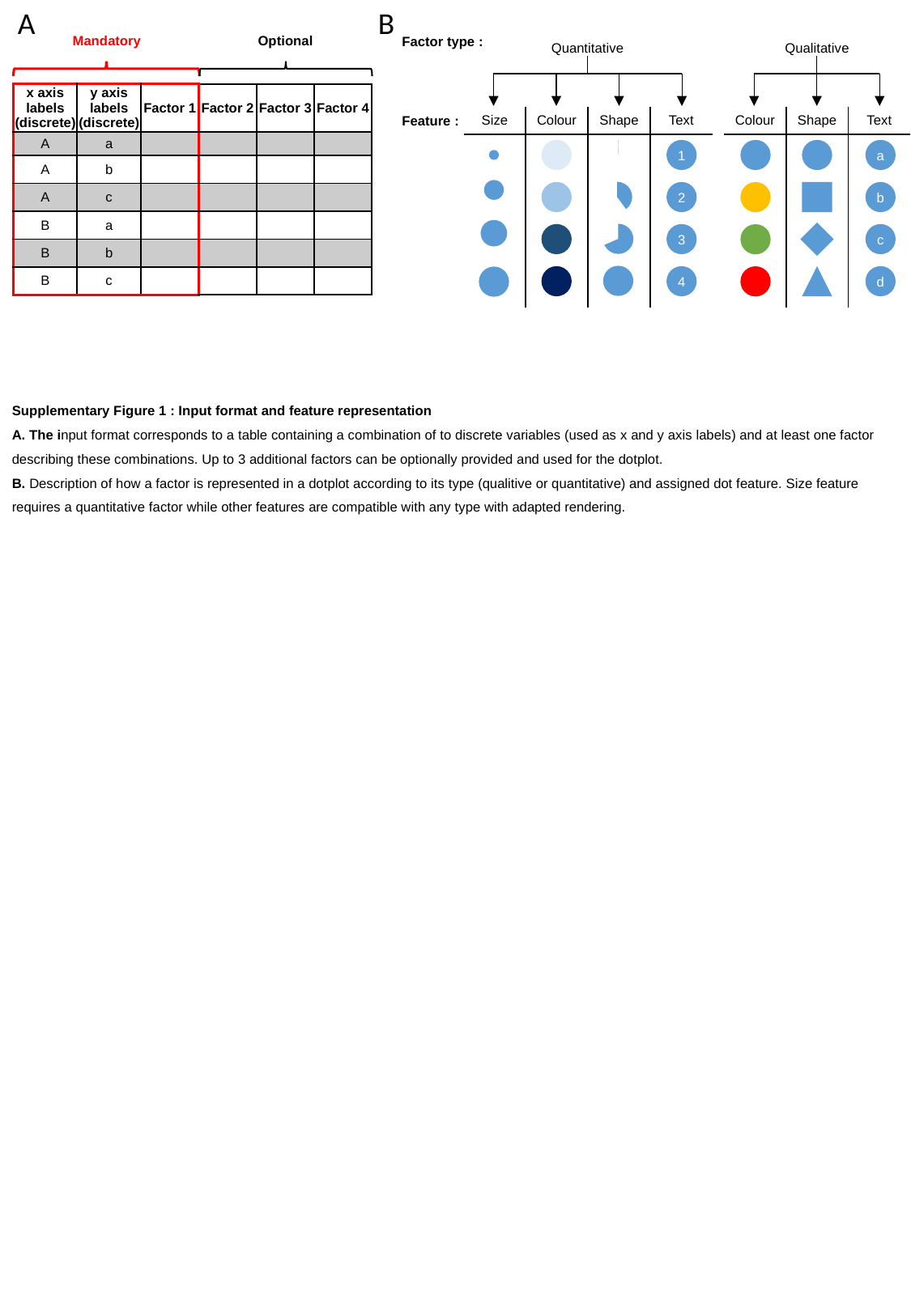

A
B
Mandatory
Optional
Factor type :
Quantitative
Qualitative
| x axis labels (discrete) | y axis labels (discrete) | Factor 1 | Factor 2 | Factor 3 | Factor 4 |
| --- | --- | --- | --- | --- | --- |
| A | a | | | | |
| A | b | | | | |
| A | c | | | | |
| B | a | | | | |
| B | b | | | | |
| B | c | | | | |
Feature :
| Size | Colour | Shape | Text |
| --- | --- | --- | --- |
| Colour | Shape | Text |
| --- | --- | --- |
1
2
3
4
a
b
c
d
Supplementary Figure 1 : Input format and feature representation
A. The input format corresponds to a table containing a combination of to discrete variables (used as x and y axis labels) and at least one factor describing these combinations. Up to 3 additional factors can be optionally provided and used for the dotplot.
B. Description of how a factor is represented in a dotplot according to its type (qualitive or quantitative) and assigned dot feature. Size feature requires a quantitative factor while other features are compatible with any type with adapted rendering.

### Slide 2
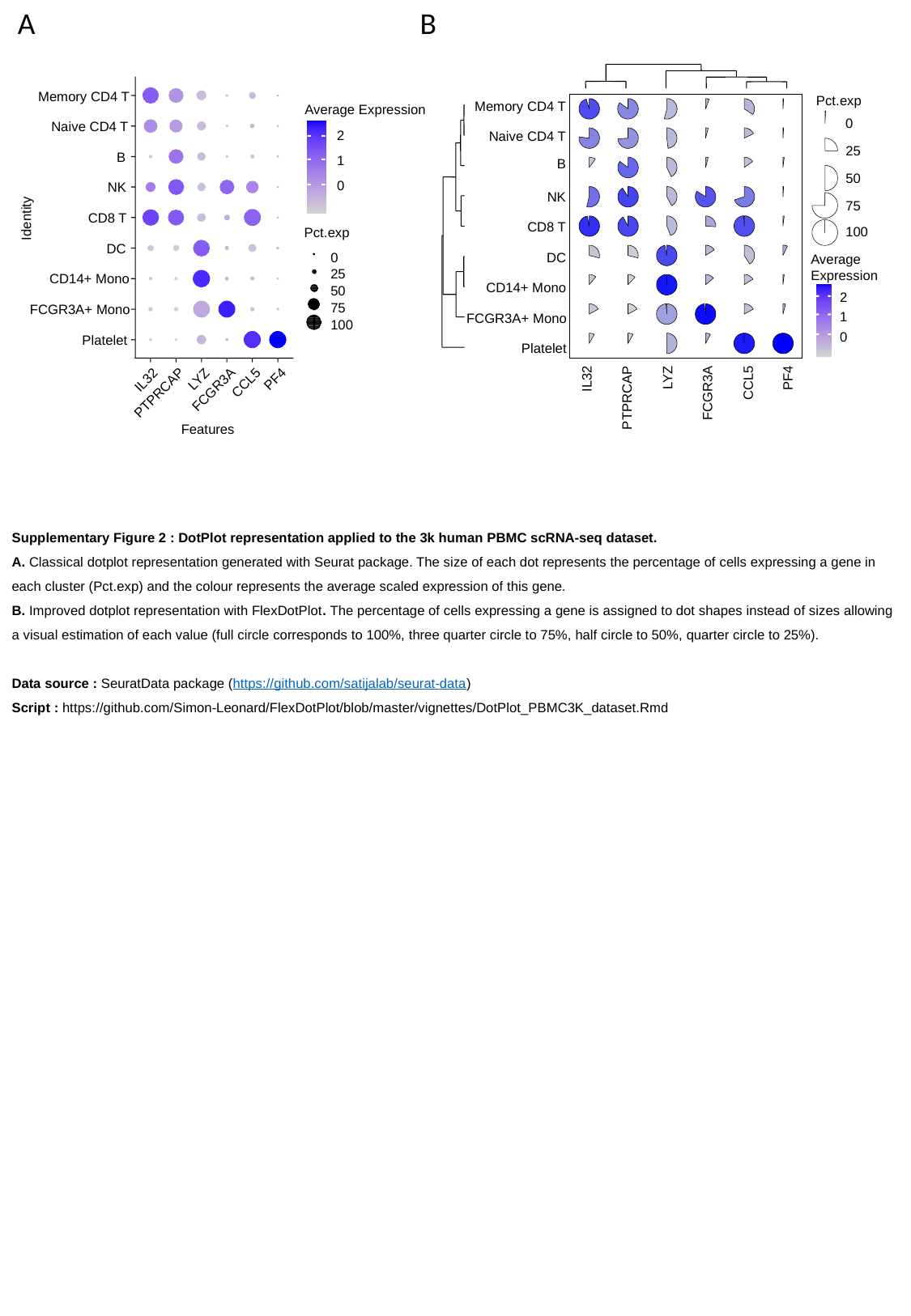

A
B
Pct.exp
Memory CD4 T
Memory CD4 T
Naive CD4 T
B
NK
CD8 T
DC
CD14+ Mono
FCGR3A+ Mono
Platelet
Average Expression
0
Naive CD4 T
2
25
B
1
50
0
NK
75
CD8 T
Identity
100
Pct.exp
DC
0
Average Expression
25
CD14+ Mono
50
2
1
0
75
FCGR3A+ Mono
100
Platelet
LYZ
PF4
IL32
CCL5
FCGR3A
PTPRCAP
IL32
PTPRCAP
LYZ
FCGR3A
CCL5
PF4
Features
Supplementary Figure 2 : DotPlot representation applied to the 3k human PBMC scRNA-seq dataset.
A. Classical dotplot representation generated with Seurat package. The size of each dot represents the percentage of cells expressing a gene in each cluster (Pct.exp) and the colour represents the average scaled expression of this gene.
B. Improved dotplot representation with FlexDotPlot. The percentage of cells expressing a gene is assigned to dot shapes instead of sizes allowing a visual estimation of each value (full circle corresponds to 100%, three quarter circle to 75%, half circle to 50%, quarter circle to 25%).
Data source : SeuratData package (https://github.com/satijalab/seurat-data)
Script : https://github.com/Simon-Leonard/FlexDotPlot/blob/master/vignettes/DotPlot_PBMC3K_dataset.Rmd

### Slide 3
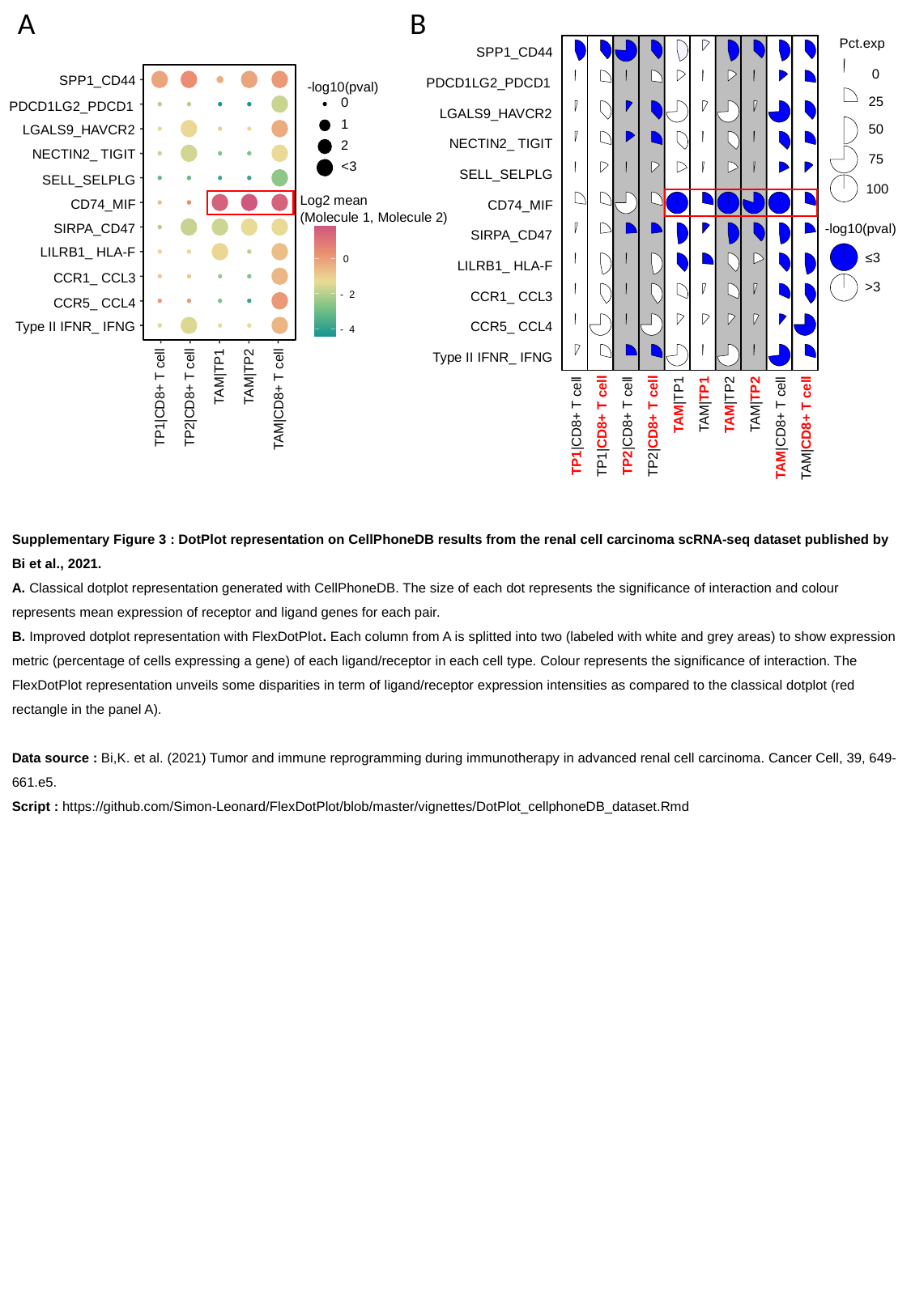

A
B
Pct.exp
SPP1_CD44
0
PDCD1LG2_PDCD1
25
LGALS9_HAVCR2
50
NECTIN2_ TIGIT
75
SELL_SELPLG
100
CD74_MIF
-log10(pval)
SIRPA_CD47
≤3
LILRB1_ HLA-F
>3
CCR1_ CCL3
CCR5_ CCL4
Type II IFNR_ IFNG
TAM|TP1
TAM|TP1
TAM|TP2
TAM|TP2
TP1|CD8+ T cell
TP1|CD8+ T cell
TP2|CD8+ T cell
TP2|CD8+ T cell
TAM|CD8+ T cell
TAM|CD8+ T cell
SPP1_CD44
TP1|CD8+ T cell
TP2|CD8+ T cell
TAM|TP1
TAM|TP2
TAM|CD8+ T cell
-log10(pval)
PDCD1LG2_PDCD1
0
LGALS9_HAVCR2
1
2
NECTIN2_ TIGIT
<3
SELL_SELPLG
CD74_MIF
Log2 mean
(Molecule 1, Molecule 2)
SIRPA_CD47
LILRB1_ HLA-F
0
CCR1_ CCL3
-
2
CCR5_ CCL4
Type II IFNR_ IFNG
-
4
Supplementary Figure 3 : DotPlot representation on CellPhoneDB results from the renal cell carcinoma scRNA-seq dataset published by Bi et al., 2021.
A. Classical dotplot representation generated with CellPhoneDB. The size of each dot represents the significance of interaction and colour represents mean expression of receptor and ligand genes for each pair.
B. Improved dotplot representation with FlexDotPlot. Each column from A is splitted into two (labeled with white and grey areas) to show expression metric (percentage of cells expressing a gene) of each ligand/receptor in each cell type. Colour represents the significance of interaction. The FlexDotPlot representation unveils some disparities in term of ligand/receptor expression intensities as compared to the classical dotplot (red rectangle in the panel A).
Data source : Bi,K. et al. (2021) Tumor and immune reprogramming during immunotherapy in advanced renal cell carcinoma. Cancer Cell, 39, 649-661.e5.
Script : https://github.com/Simon-Leonard/FlexDotPlot/blob/master/vignettes/DotPlot_cellphoneDB_dataset.Rmd

### Slide 4
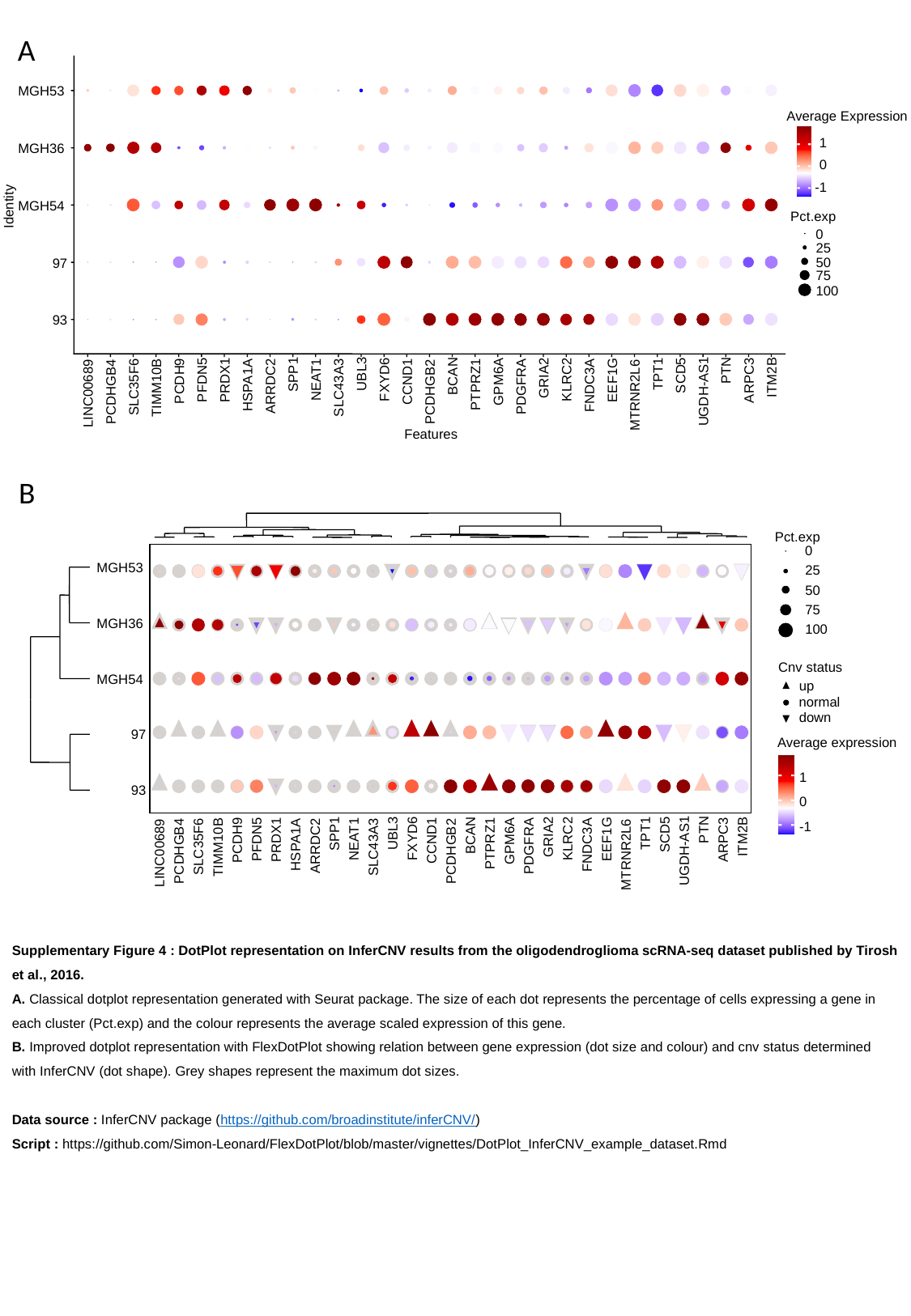

A
MGH53
Average Expression
1
MGH36
0
-1
MGH54
Identity
Pct.exp
0
25
50
97
75
100
93
PTN
TPT1
UBL3
SPP1
SCD5
BCAN
ITM2B
GRIA2
FXYD6
KLRC2
NEAT1
PFDN5
EEF1G
PRDX1
ARPC3
PCDH9
CCND1
GPM6A
PTPRZ1
HSPA1A
FNDC3A
ARRDC2
PDGFRA
SLC35F6
SLC43A3
TIMM10B
UGDH-AS1
PCDHGB4
PCDHGB2
LINC00689
MTRNR2L6
Features
B
Pct.exp
0
MGH53
25
50
75
MGH36
100
Cnv status
MGH54
up
normal
down
97
Average expression
1
93
0
-1
PTN
TPT1
UBL3
SPP1
SCD5
BCAN
ITM2B
GRIA2
FXYD6
KLRC2
NEAT1
PFDN5
EEF1G
PRDX1
ARPC3
PCDH9
CCND1
GPM6A
PTPRZ1
HSPA1A
FNDC3A
ARRDC2
PDGFRA
SLC35F6
SLC43A3
TIMM10B
UGDH-AS1
PCDHGB4
PCDHGB2
LINC00689
MTRNR2L6
Supplementary Figure 4 : DotPlot representation on InferCNV results from the oligodendroglioma scRNA-seq dataset published by Tirosh et al., 2016.
A. Classical dotplot representation generated with Seurat package. The size of each dot represents the percentage of cells expressing a gene in each cluster (Pct.exp) and the colour represents the average scaled expression of this gene.
B. Improved dotplot representation with FlexDotPlot showing relation between gene expression (dot size and colour) and cnv status determined with InferCNV (dot shape). Grey shapes represent the maximum dot sizes.
Data source : InferCNV package (https://github.com/broadinstitute/inferCNV/)
Script : https://github.com/Simon-Leonard/FlexDotPlot/blob/master/vignettes/DotPlot_InferCNV_example_dataset.Rmd
